# Brownian motion data augmentation: a method to push neural network performance on nanopore sensors

**DOI:** 10.1101/2024.09.10.612270

**Authors:** Javier Kipen, Joakim Jaldén

## Abstract

Nanopores are highly sensitive sensors that have achieved commercial success in DNA/RNA sequencing, with potential applications in protein sequencing and biomarker identification. Solid-state nanopores, in particular, face challenges such as instability and low signal-to-noise ratios (SNRs), which lead scientists to adopt data-driven methods for nanopore signal analysis, although data acquisition remains restrictive. In this paper, we augment training samples by simulating virtual Brownian motion based on dynamic models in the literature. We apply this method to a publicly available dataset of a classification task containing nanopore reads of DNA with encoded barcodes. A neural network named QuipuNet was previously published for this dataset, and we demonstrate that our augmentation method produces a noticeable increase in QuipuNet’s accuracy. Furthermore, we introduce a novel neural network named YupanaNet, which achieves greater accuracy (95.8%) than QuipuNet (94.6%) on the same dataset. YupanaNet benefits from both the enhanced generalization provided by Brownian motion data augmentation and the incorporation of novel architectures, including skip connections and a self-attention mechanism.

## 2 Introduction

Nanopore sensors operate by measuring a current of ions passing through a nanometer-sized pore when a voltage is applied across the membrane. Any molecule traversing the pore modulates this ion current, allowing for identification based on the modulation pattern. Oxford Nanopore Technologies stands as a prominent example of commercial success in this domain, particularly in DNA sequencing [1]. However, the potential applications of nanopore sensors extend beyond DNA sequencing, encompassing fields such as protein sequencing [2, 3, 4, 5, 6], biomarker identification [7], and DNA-based data storage [8, 9]. Despite the expanding scope of this field, data acquisition and processing remain a limiting factor.

A specific sub-field gaining traction is DNA-based nanopore sensing [10, 11], which involves translocating DNA through pores while concurrently measuring other components attached to the DNA molecule. DNA can translocate easily through nanopores since it is electrically charged, which enables binding uncharged targets to the DNA and measuring both. Applications include structural data storage on DNA [12, 13, 14, 15], single-molecule detection [16, 17, 18], and binding site identification [19, 20].

Signal processing methods can be applied to analyze DNA-based nanopore reads originating from solid-state nanopores [21, 22, 23, 24]. However, solid-state nanopores are often unstable and exhibit substantial variations, necessitating extensive algorithm parameter tuning. This task is particularly hard within DNA-based sensing, since there is an intrinsically lower signal-to-noise ratio. Data-driven approaches can achieve better results with less time spent on data analysis.

One illustrative example is the case of digitally multiplexing reads of nanopore [18]. In the mentioned paper the authors introduce hairpins to encode a digital barcode that indicates what is to be tested and an antibody on the other side of the DNA that binds to a specific molecule. Although they achieved a high accuracy (94%) in the barcode detection, events where there is DNA folding were discarded. In a following study Misiunas et al. [25], the authors introduced a Convolutional Neural Network (CNN), and they showed that it can achieve higher accuracies when considering a significantly bigger (5-fold bigger) event dataset containing DNA folds. Despite there being other examples of neural networks applied to similar data [26, 27], the field remains relatively unexplored, and there are not many examples of expert knowledge of this particular field applied to boost data-driven methods performance.

Concurrently, there are studies analyzing the translocation of DNA through a nanopore both via simulations and experimentally [28, 29, 30]. In these works, the impact of thermal forces on translocation dynamics were described. At the same time, data augmentation techniques have demonstrated efficacy in enhancing neural network performance in time series classification tasks [31], particularly in domains with limited data availability, which is the case for nanopore sensing. Therefore, we introduce a novel data augmentation method termed “Brownian Motion Data Augmentation,” which emulates the effect of thermal forces on the real samples. We showcase in our results an improvement in accuracy for the existing state-of-the-art neural network provided by Misiunas et al. [25].

Additionally, we fine-tune a new neural network architecture leveraging the benefits of Brownian motion data augmentation to achieve even higher accuracies. Given the success of residual networks [32] and attention mechanisms [33] in the area of deep learning, we proposed a neural network based on the existing QuipuNet [25] adding skip-connections and self-attention blocks. Our proposed model achieves a top test accuracy of 95.8% compared to the previously reported 94.6% of QuipuNet. Since the “Quipu” was an Incan mathematical tool, we decided to continue referencing the Incan empire mathematical tools and named our neural network YupanaNet.

## 3 Methods

### 3.1 QuipuNet summary

QuipuNet, introduced by Misiunas et al. [25], is a Convolutional Neural Network designed to classify both barcodes and the presence of a target molecule in DNA-based reads from solid-state nanopores. These recorded traces are ideally a drop in the ion current due to the DNA translocating, and other drops, along the previous one, which represent the barcode and the presence (or not) of a target molecule. The barcode signal consists of two indicating dips, between which there may or may not be up to three intermediate dips, with the presence of a drop indicating a binary 1, and the absence a binary 0. Consequently, the barcode identification is a classification task with eight classes, the number of possible classes indexed by three bits. It is worth mentioning that DNA translocation can occur in both directions within the traces, resulting in variability in the position of barcodes, and traces may exhibit additional peaks due to DNA folding. Although Quipu was designed for both barcode and target detection, we focus on the barcode detection in this paper.

The dataset introduced in [25] comprises 55,989 samples, with 52,525 allocated for training. Notably, the eight barcodes within the dataset vary considerably in sample count and the traces can have a target molecule bounded or not. The authors preprocessed the dataset by removing events likely stemming from contamination, erroneous detection, and incomplete DNA fragments. In addition, they separated traces according to which experiment they belonged to, to ensure that the events on the training data were not from the same experiment as the ones in the test set. This separation aided in training the neural network to classify based on trace characteristics rather than experimental factors, preventing it from identifying experiments based on pore characteristics. Additionally, current values were normalized relative to the experiment’s unfolded current level. During neural network training, the dataset was augmented in each epoch by linearly stretching and squeezing samples, adding noise, and applying baseline variations to the entire trace. Detailed parameter values and configurations for training are available in their GitHub repository [34].

### 3.2 Brief summary of dynamics of DNA translocations on nanopores

The translocation of DNA through the pore is described in the literature [29, 30] by the one-dimensional Langevin equation

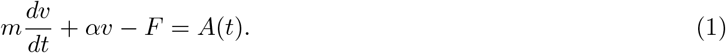

Here, *m* represents the mass of the DNA molecule, *F* = *V*_b_*λ* denotes the driving electrical field force, with *λ* being the charge density of the DNA and *V*_b_ is the bias applied. The term *A*(*t*) accounts for stochastic thermal forces caused by random molecular collisions, and *αv* constitutes the drag force opposing the relative motion of the DNA.

Due to its simplicity, (1) doesn’t precisely align with experimental data. To refine this model, considerations of the dynamic change in the drag coefficient during DNA translocation through the nanopore are necessary. As explored in [29], modeling DNA as an unraveling chain simplifies drag coefficient calculations and accurately approximates the physical scenario. Additionally, introducing a pivot point, beyond which DNA segments have negligible movement, simplifies computations while preserving accurate modeling. These model refinements are considered when rewriting the Langevin equation as

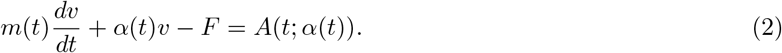

In (2), *m*(*t*) is the mass of the DNA before the pivot point, the part that is being untangled and translocated, and the drag coefficient *α*(*t*) is calculated considering each chain segment. This adjusted model has been validated against experimental data [29], demonstrating its capability to represent observed translocation dynamics accurately.

One significant insight from simulations, as highlighted in [29], is the impact of thermal noise on translocation speed. Incorporating thermal forces introduces additional noise, resembling white noise, into the instantaneous translocation speeds. This finding serves as the foundation for modeling our Brownian motion augmentation.

### 3.3 Brownian motion data augmentation

We introduce Brownian motion data augmentation as a method to create a virtual trace *Y* [*n*] ∈ ℝ^*L*^ from a real trace *X*[*n*] ∈ ℝ^*L*^ by simulating other possible thermal force interactions than those that were in effect when the trace was measured. The length *L* accounts for the number of samples of the trace. As mentioned in the previous subsection, an uncorrelated noise is added to the instantaneous translocation speed when thermal forces are considered in the simulations. Although it is a simplification, we model the noise as Gaussian. Consequently, the stochastic process of DNA’s instantaneous discrete position as it translocates through the nanopore can be modeled as a summation of its discrete instantaneous speed, i.e., a biased Brownian random walk.

One simplification used in our method is to estimate the DNA instantaneous position directly from the measured current. The nanopore capacitance and solution resistance act as a filter, but in order to model them these parameters would need to be estimated for every experiment. In addition, we model the thermal forces as being stationary during the whole translocation instead of them depending on time as in (2). This simplification allows us to do the augmentation without the details of the unraveling of the DNA, which would be cumbersome to estimate only from the traces. Despite these simplifications, our method can still boost the neural networks’ performance in the barcode classification task.

After mentioning the simplifications, we are now able to introduce the augmentation method. The first step in our method involves obtaining newly generated output indexes *i*_0_[*n*], which indicate the new virtual instantaneous positions of the DNA. In the absence of thermal noise, these indexes are natural numbers up to *L*. When considering thermal noise, the index is summed with a random variable *Z*[*n*] ∼ 𝒩(1, *σ*). *Z*[*n*] represents the position advancement has a mean of one sample and added uncorrelated Gaussian noise, modelling the effect of the thermal noise on the instantaneous position. The resulting expression is

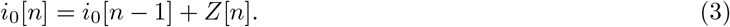

It’s important to note that with increasing time, the variance of the index increases with 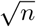, consistent with the behavior of a Brownian motion process. However, indexes may become negative and this is solved in practice by replacing the negative numbers with zeroes.

Once the new indexes are obtained, the augmented trace *Y* [*n*] is obtained by interpolating from the original trace *X*[*n*]. Defining *X*_⌊_[*n*] = *X*[⌊*i*_0_[*n*]⌋] and *X*_⌈_[*n*] = *X*[⌈*i*_0_[*n*]⌉], we linearly interpolate from *X*[*n*] to get *Y* [*n*] as

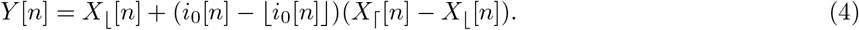

This interpolation approach is straightforward and amenable to efficient parallelization. Using other interpolation methods, such as splines or Fourier resampling, could result in a more accurate reconstruction. However, for the dataset used, the index variation due to this augmentation is relatively small, and this simple implementation could already show noticeable accuracy improvements.

An example of practical augmentation is plotted in Figure 1. First, the augmenting indexes are obtained using (3) and *σ* = 0.9, and they are shown in Figure 1a. For this example *L* = 700 since the event data was padded as in [25] to standardize the dataset trace length. Notably, the resulting augmenting indexes *i*_*o*_[*n*] indicate that the process achieves some latter positions earlier, suggesting faster translocation at the beginning, followed by reduced speed. In the interpolated signal in Figure 1b these speed variations result in the first peak being forwarded, and other peaks being delayed. Figure 1b demonstrates an effect of both squeezing and stretching the signal in the same trace.

**Figure 1:**
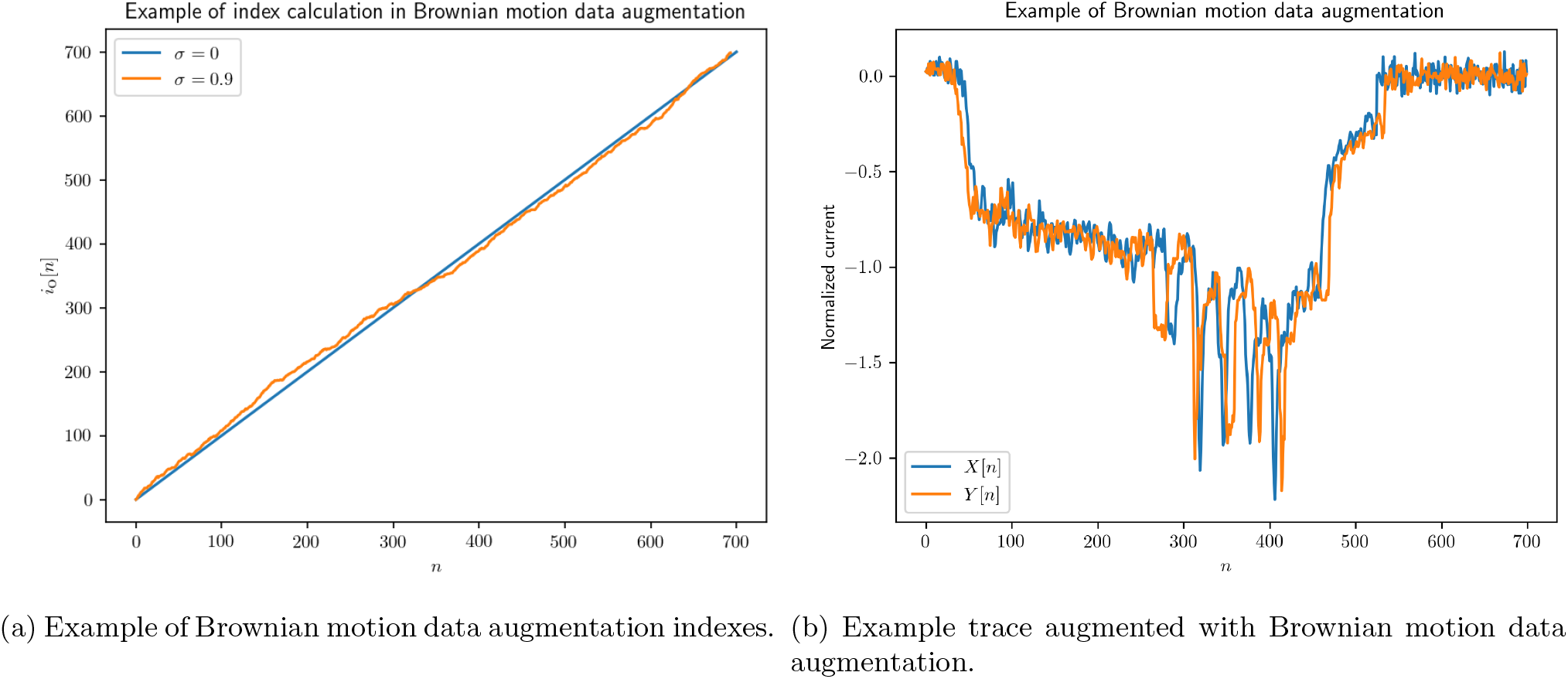
Example of the Brownian motion data augmentation method

### 3.4 YupanaNet

Besides introducing Brownian motion augmentation, we also propose a novel neural network architecture, YupanaNet, to further improve the accuracy of the barcode classification task. This experiment also allows us to test if our proposed data augmentation allows us to train more complex neural networks without overfitting. Inspired by recent advancements in neural network architectures, YupanaNet incorporates both residual connections and an attention mechanism. These architectural elements have shown promising results in various time series classification tasks.

While Convolutional Neural Networks (CNNs) have been widely used for time series classification problems, recent studies [35] suggest that residual neural networks often outperform them in those problems. ResNets, initially introduced for image classification [32], have demonstrated the ability to train deeper models and achieve higher accuracies.

In parallel, transformer-based architectures, known for their success in natural language processing (NLP) tasks, have also gained attention. Transformers, introduced by Vaswani et al. [33], leverage self-attention mechanisms to enable the network to focus on relevant features within the input sequence. This attention mechanism has proven effective in capturing long-range dependencies, making transformers a promising choice for sequence modeling tasks, including time series analysis [36].

Given the success of both ResNets and transformers, along with the original QuipuNet’s state-of-the-art performance, we incorporate residual connections and an attention module into the QuipuNet architecture. Similar combinations have been applied in various time series classification tasks in the literature [37, 38, 39]. Additionally, we expanded the network architecture by adding more layers and increasing the size of some layers.

The final model and hyper-parameters were carefully tuned, with further descriptive details provided in the supplementary information. The architecture of YupanaNet is illustrated in Figure 2a. It begins with a convolutional layer followed by a self-attention block. Subsequently, it comprises a series of blocks inspired by the QuipuNet structure but incorporating residual connections. Finally, a flatten layer is followed by two fully connected layers.

**Figure 2:**
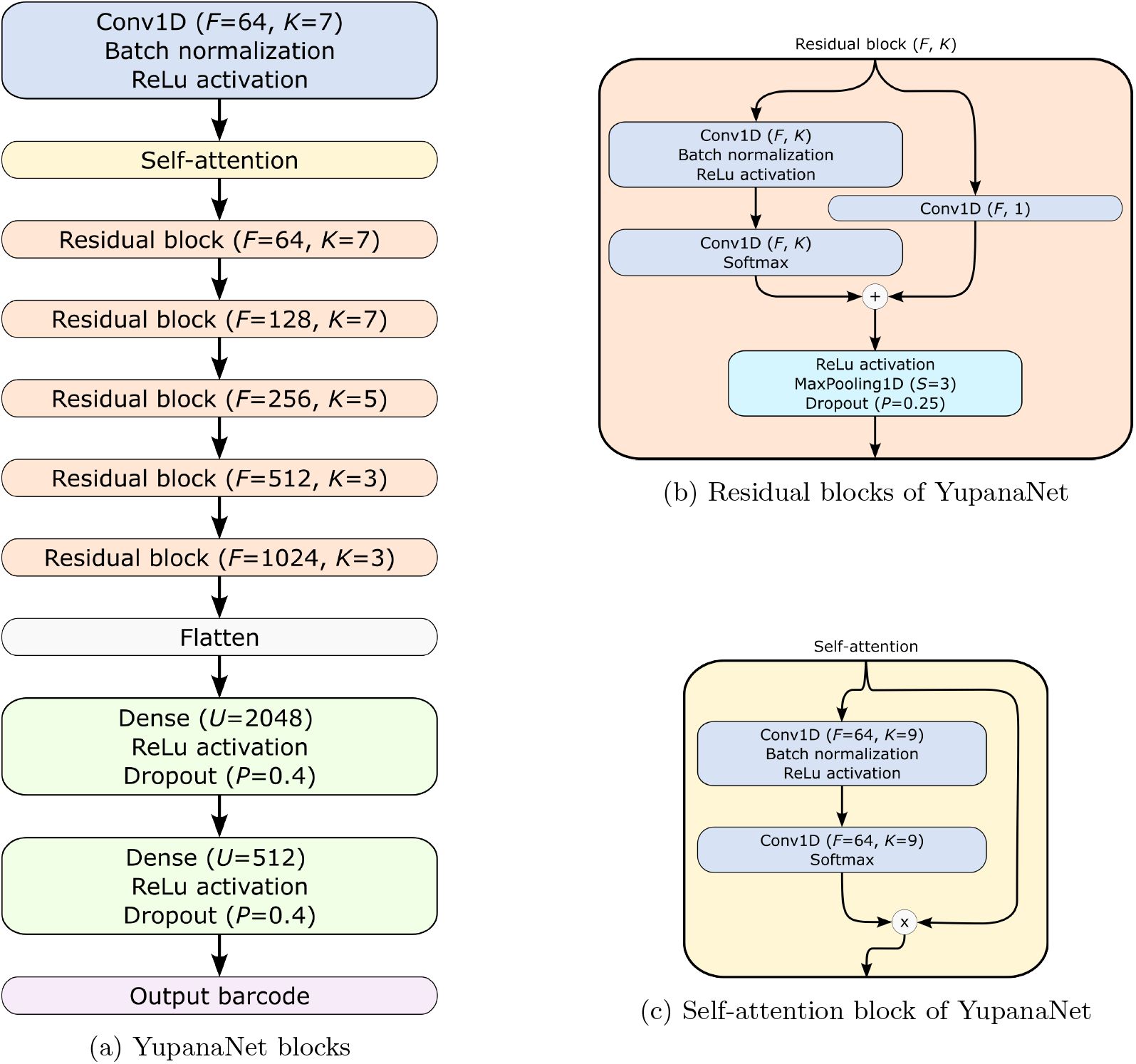
YupanaNet architecture. Here, *F* denotes the number of filters in each convolutional layer, *K* represents the size of the convolution window, *U* is the number of units in the dense layer, *P* indicates the dropout probability, and *S* specifies the size of the max pooling window.

The residual connection blocks, depicted in Figure 2b, maintain the same structure as the convolutional blocks of QuipuNet but include a skip connection facilitated by passing through a 1D convolutional filter with a kernel size of 1. The self-attention block, shown in Figure 2c, incorporates the attention mechanism, allowing the network to selectively focus on relevant features during processing.

## 4 Results

This section comprises two main parts, each focusing on different aspects while utilizing the Quipu dataset [25]. In the first part, we demonstrate the efficacy of Brownian augmentation in enhancing the performance of QuipuNet. In the second part, we demonstrate the accuracy gain obtained gained with YupanaNet. The training of the Neural Networks was done on the Alvis cluster, an NVIDIA-based cluster with focus on Artificial Inteligence and Machine Learning research, provided by The National Academic Infrastructure for Supercomputing in Sweden (NAISS).

### 4.1 Improvement of accuracy due to Brownian motion data augmentation

#### 4.1.1 Dataset and training

For this part of the results, we generated random training and test datasets from the complete dataset of Misiunas et al. [25] to better represent generalization accuracy. Our objective here was to conduct multiple random runs to demonstrate that the accuracy increase observed with our augmentation was not merely due to randomness.

Although the test set was randomly selected, we maintained the constraint of using traces from different experiments for the training and test. The randomization process followed several steps: first, for each barcode (label), we determined the percentage of traces of each experiment. Additionally, we compiled a list of all possible test sets that could be generated with the mentioned constraint and that the test set for each label represented between 4% and 15% of the traces from the same label. Finally, a combination of those is selected at random and the test set is generated when applying these steps to all labels. This process ensured picking random test sets that were from different experiments compared to the training dataset.

We initially set aside 10% of the training samples as a validation set, ensuring that the validation set maintained the same class proportions as the training set. The resulting training dataset was oversampled to balance out the classes, and each sample was augmented every epoch. The augmentation process aligned with the parameters used in Quipu training, including independently stretching, noise addition, and magnitude augmentation. We compared training with and without our Brownian motion augmentation technique with a fixed reasonable parameter (*σ* = 0.9). Additionally, since the use of Brownian augmentation resulted in traces of varying lengths, we also compared it with a stretching method that introduced the same length variation as the Brownian augmentation. This step ensured that any observed increase in accuracy was not solely due to the varying length caused by the Brownian augmentation. Training proceeded for 100 epochs, and the model weights with the best validation loss were used to evaluate the test set. The batch size was fixed to 256, and different learning rate (LR) values were tested.

#### 4.1.2 Accuracy results

We compared the accuracy with the QuipuNet architecture and an extended version in which we added more nodes to the two last dense layers (from [512, 512] to [2048, 1024]). The trainings were run with and without Brownian augmentation and also varying the learning rates, and all the results are shown in the Supplementary Information. Finally, we pick the best test accuracy results for each architecture and augmentation configuration, and the results are shown in Table 1. The accuracies are presented as mean values ± the standard deviation of the mean accuracy estimator, while the last column represents the 90% percentile.

**Table 1:** Accuracy comparison.

| $N$ Dense | Brow | Stretch eq | LR | Train acc | Test acc | Test acc best 90% |
| --- | --- | --- | --- | --- | --- | --- |
| [512,512] | No | No | 1E-03 | $99.3 \pm 0.01\%$ | $92.95 \pm 0.11\%$ | 95.1% |
| [512,512] | **Yes** | No | 1E-03 | $98.18 \pm 0.01\%$ | $93.12 \pm 0.11\%$ | 95.1% |
| [2048,1024] | No | No | 5E-04 | $99.14 \pm 0.02\%$ | $93.04 \pm 0.11\%$ | 95.2% |
| [2048,1024] | **Yes** | No | 5E-04 | $98.67 \pm 0.01\%$ | **$93.42 \pm 0.10\%$** | **95.7%** |
| [2048,1024] | No | Yes | 5E-04 | $99.01 \pm 0.01\%$ | $93.03 \pm 0.09\%$ | 95.3% |

We could reproduce even higher accuracies than those reported for QuipuNet [25] (94.6%), with a 90% percentile of 95.1%, but this increase is because other test datasets were used. We can also appreciate that the mean test accuracy with Brownian augmentation on QuipuNet is higher but not considerably. While this test accuracy shows improvement, it’s important to note a significant drop in training accuracy, suggesting increased complexity of the classification task for the neural network when using the augmentation, which in turn reduces overfitting.

The best test accuracies of the extended model with and without Brownian motion data augmentation are shown as the third and fourth row in Table 1. These rows show that enlarging the architecture with the same augmentations didn’t result in a noticeable change in the test accuracy. However, when using Brownian motion data augmentation, both train and test mean accuracy increased compared to the smaller model. We interpret that this bigger neural network can solve the more complex classification task when using Brownian augmentation. Considering the variances of the mean accuracy estimators, this increase in accuracy is unlikely to be attributed to randomness. Additional calculations are included in the supporting information to support this conclusion.

Finally, we trained the larger neural network, replacing Brownian motion augmentation with uniform stretching/squeezing equivalent to the one produced by our augmentation. The equivalent stretch is obtained by observing the variance in the relative length of the events due to our augmentation method, and the equivalent stretch empirical estimation is described in the Supporting information. The results, shown in the last row of Table 1, indicated no increase in accuracy. This result confirms that the accuracy enhancement in the extended model was indeed due to Brownian motion augmentation.

### 4.2 Results of YupanaNet

#### 4.2.1 Dataset and training

For this part of the results, we used exactly the same training and test dataset as QuipuNet [25]. The test set was separated while tuning YupanaNet, assuring no test overfitting. A more detailed description of how YupanaNet was tuned can be found on the supplementary information.

A validation set was separated from the training dataset with 5% of the samples, assuring also same percentage for each class. Additionally, the remaining train dataset was oversampled to ensure that each class had the same amount of samples. During the training, the training dataset was augmented independently for each “epoch” with stretching, noise addition, magnitude multiplication and Brownian augmentation. In addition, these augmentations were optimized in computing time so that their running time became negligible in the training. The parameters of the augmentations were also tuned. Finally, the training consisted of fitting 100 epochs with cross-entropy loss and picking the model weights which minimized the validation dataset loss. The chosen optimizer was Adam [40], with a tuned learning rate and batch size.

#### 4.2.2 Accuracy comparison

Table 2 compares the test accuracies of different published models and YupanaNet. The test’s accuracy is shown as the mean ± the standard deviation of tens of runs on the same training dataset. Results of a previously published alternative models, T-S2Inet [41] and S2Snet [42], were not included in the comparison because their training could not be reproduced with the provided GitHub repositories

**Table 2:**
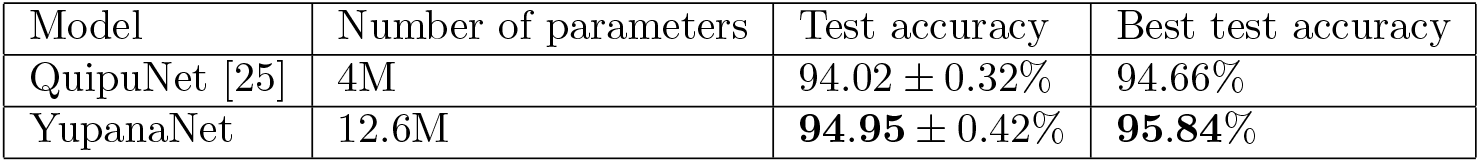
Accuracy comparison.

QuipuNet was trained 117 times and a slightly higher accuracy than the reported in the paper was achieved. Table 2 shows that YupanaNet achieves higher mean and best test accuracy than QuipuNet and S2SNET. This increase is due to the Brownian motion data augmentation and the newer architecture of YupanaNet.

#### 4.2.3 Ablation study

We finally performed an ablation study on YupanaNet to show the contribution of both the Brownian motion data augmentation and the self-attention module. The residual connections were not studied since there is significant literature which justifies that adding residual connections improves the results of these neural networks. The ablation results are shown in Table 3, where each configuration was run approximately 30 times.

**Table 3:** Ablation studies table.

| Brownian DA | Self-Attention | Test accuracy | Best test accuracy |
| --- | --- | --- | --- |
| Yes | Yes | **94.95 ± 0.44%** | **95.84%** |
| Yes | No | 94.79 ± 0.34% | 95.41% |
| No | Yes | 94.57 ± 0.45% | 95.24% |
| No | No | 94.44 ± 0.29% | 94.95% |

Table 3 shows that both modules help achieve the highest accuracy, and whenever one is removed, the accuracy drops. In addition, when removing both modules results are very similar to QuipuNet, although a slightly higher accuracy is achieved. This increment can be explained by the skip connections added to the structure and its larger size. This study shows that both the Brownian motion data augmentation and the self-attention block significantly contribute to the increase in the accuracy of YupanaNet.

## 5 Discussion

This paper introduces two significant contributions: the Brownian motion data augmentation method and YupanaNet, a novel neural network architecture with residual connections and a self-attention block. The experiments demonstrate the efficacy of the Brownian motion augmentation in enhancing the accuracy of QuipuNet, and that YupanaNet outperforms existing state-of-the-art models.

The presented Brownian motion augmentation method, while simple, showcases enhanced results in the mentioned barcode classification task. Although further refinements could consider factors like the filtering effect of the nanopore’s capacitance and more accurate noise models of the instantaneous velocity due to thermal forces, this method presents a viable and accessible means of enhancing neural network performance in DNA-based nanopore sensing. While the observed accuracy improvement may not be huge, its ease of implementation makes it a valuable tool, particularly in light of the ongoing challenges associated with nanopore data collection.

Moreover, the Brownian motion augmentation method holds promise for applications beyond DNA-based nanopore sequencing. Since it results from thermal forces applied to molecules translocating through a pore, it could potentially be practical for other nanopore measurements that do not rely on motor proteins. For instance, it could be applicable to augment data for data-driven methods applied translocating proteins [2, 5, 6, 5]. The simplicity and effectiveness of this method suggest its potential applicability in various physical systems where Brownian motion affects the measurements. Thus, its utility extends beyond nanopore sensing.

In parallel, YupanaNet represents a notable advancement in nanopore classification tasks. By incorporating residual connections and a self-attention mechanism, YupanaNet builds upon the success of existing neural network architectures while offering improved accuracy and performance in the barcode classification task of Misiunas et al. [25]. Its success positions YupanaNet as a noteworthy example for similar classification tasks within the field. Given the common occurrence of variable length inputs in nanopore data analysis, exploring the integration of global averaging layers to handle such inputs, or investigating the implementation of recursive structures, could hold promise for advancing the capabilities of neural networks in nanopore-based classification tasks.

## 6 Acknowledgements

This work has been supported by the Swedish Research Council (VR Research Environment Grant 2018-06169; QuantumSense), and the Swedish Foundation for Strategic Research (SSF Grant ITM17-0049). The computations were enabled by resources provided by the National Academic Infrastructure for Supercomputing in Sweden (NAISS), partially funded by the Swedish Research Council through grant agreement no. 2022-06725.

ChatGPT was used to improve the cohesion, style, and vocabulary of the text, and Github Copilot was used for the development of the code.

---

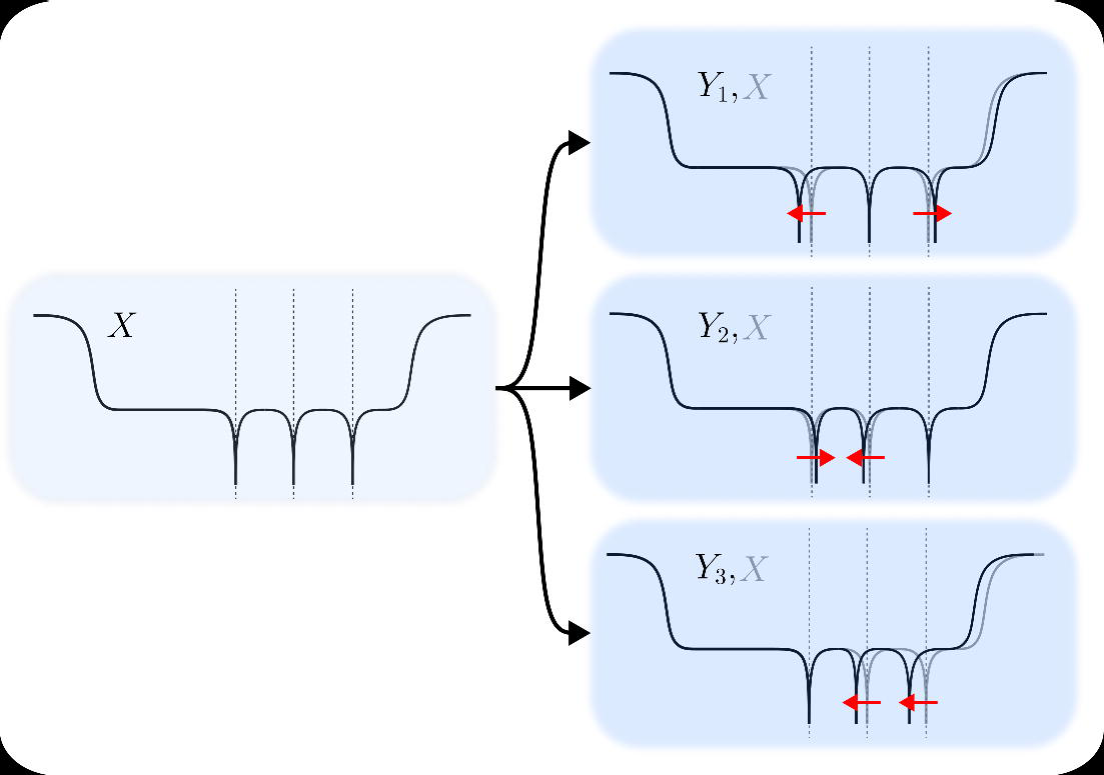

## Notes

https://github.com/kmisiunas/QuipuNet

### Competing Interest Statement

The authors have declared no competing interest.

