## Supplementary information for "Brownian motion data augmentation: a method to push neural network performance on nanopore sensors"

Javier Kipen, Joakim Jalden

August 2024

### 1 Supporting information

#### 1.1 Results of learning rate sweep in the neural networks trainings

All the results of the trainings for different learning rates (LR) and different configurations of the neural networks are shown in Table 1.

| N Dense | Brow | Stretch eq | LR | Train acc | Test acc | N runs |
|---|---|---|---|---|---|---|
| [512,512] | No | No | 5E-3 | $95.91 \pm 1.83\%$ | $88.98 \pm 1.83(93.97)\%$ | 54 |
| [512,512] | No | No | 2E-3 | $99.27 \pm 0.01\%$ | $92.86 \pm 0.16(94.91)\%$ | 108 |
| [512,512] | No | No | 1E-3 | $99.35 \pm 0.01\%$ | $92.96 \pm 0.12(95.11)\%$ | 208 |
| [512,512] | No | No | 5E-4 | $99.16 \pm 0.02\%$ | $92.76 \pm 0.14(94.79)\%$ | 150 |
| [512,512] | No | No | 3E-4 | $98.80 \pm 0.03\%$ | $92.82 \pm 0.13(94.94)\%$ | 148 |
| [2048,1024] | No | No | 2E-3 | $99.32 \pm 0.02\%$ | $92.45 \pm 0.16(95.07)\%$ | 159 |
| [2048,1024] | No | No | 1E-3 | $99.22 \pm 0.02\%$ | $92.85 \pm 0.13(95.14)\%$ | 212 |
| [2048,1024] | No | No | 5E-4 | $99.15 \pm 0.02\%$ | $93.04 \pm 0.11(95.24)\%$ | 260 |
| [2048,1024] | No | No | 2E-4 | $99.07 \pm 0.03\%$ | $93.00 \pm 0.17(95.30)\%$ | 114 |
| [512,512] | Yes | No | 1E-3 | $98.18 \pm 0.01\%$ | $93.12 \pm 0.12(95.22)\%$ | 235 |
| [512,512] | Yes | No | 5E-4 | $98.00 \pm 0.02\%$ | $93.22 \pm 0.30(95.34)\%$ | 51 |
| [2048,1024] | Yes | No | 1E-3 | $98.67 \pm 0.02\%$ | $92.94 \pm 0.20(94.75)\%$ | 68 |
| [2048,1024] | Yes | No | 5E-4 | $98.68 \pm 0.01\%$ | $93.43 \pm 0.10(95.69)\%$ | 349 |
| [2048,1024] | No | Yes | 5E-4 | $99.01 \pm 0.01\%$ | $93.04 \pm 0.09(95.27)\%$ | 395 |

Table 1: Learning rate sweep for different configurations of neural networks.

We tested several learning rates for the trainings without Brownian augmentation, both with higher and equal numbers of neurons in the last dense layers. The learning rates tested with Brownian augmentation were fewer since the aim was to demonstrate that a single configuration can achieve a higher accuracy. Finally, the stretch equivalent was only tested on the same configuration where Brownian augmentation demonstrated the highest test accuracy increase.

#### 1.2 Tests to show accuracy increase

In this section, we present two different tests to verify that the proposed Brownian motion augmentation method increased the mean test accuracy. The results for the non-augmented runs correspond to the settings in the 8th row of Table 1, and the results for the augmented runs correspond to the settings in the 13th row of the same table. We define $N$ as the number of runs used.

##### 1.2.1 Probability of higher mean calculation

The first test consists of calculating the probability that the mean test accuracy is higher when trained with Brownian motion data augmentation, given the training results.

We define the random variable $T$ as the test accuracy for a run without Brownian motion augmentation and $T_{\mathrm{a}}$ as the test accuracy when training with the augmentation method. Additionally, we introduce the random variable for their difference: $D = T_{\mathrm{a}} - T$, and denote $D_i$ for the $i$th training of the networks. Our mean estimator of the difference is defined by $\hat{\mu}_D = \frac{1}{N}\sum_{i=1}^{N} D_i$.

We rewrite the probability that the mean test accuracy of the augmented method is higher than that of the non-augmented method as

$$P(\mu_{T_a} > \mu_T | D_1, D_2 \dots D_N) = P(\mu_{T_a} - \mu_T > 0 | D_1, D_2 \dots D_N) = P(\mu_D > 0 | D_1, D_2 \cdots D_N).$$

Using the estimator $\hat{\mu}_D$ we can approximate the previous expression as $P(\hat{\mu}_D > 0)$. The distribution of $\hat{\mu}_D$ can be approximated to a normal distribution due to the central limit theorem. Since $\hat{\sigma}^2_{\hat{\mu}_D} = \frac{\hat{\sigma}^2_D}{N}$, we obtain that

$$P(\mu_{T_a} > \mu_T | D_1, D_2 \dots D_N) = P(\hat{\mu}_D > 0) \simeq 1 - \Phi\left(-\frac{\hat{\mu}_D}{\hat{\sigma}_{\hat{\mu}_D}}\right)$$

where $\Phi$ represents the cumulative function of a standard normal distribution. After computing this expression with the results, we obtain a probability of 98.75% that the mean test accuracy increases when using our proposed data augmentation method.

#### 1.2.2 P-value test

The second test involves rejecting the hypothesis that there is no improvement when training with Brownian motion data augmentation. We define our null thesis ($H_0$) as the Brownian motion augmentation does not improve the mean test accuracy. Subsequently, we derive the mathematical expression to be able to calculate the probability of the observations under the null hypothesis, and the low resulting probability allows us to reject the null hypothesis.

We introduce the binary random variable $T_i$, which represents whether the test accuracy was higher with Brownian augmentation than without it on the $i$th run. $T_i$ takes the value 1 when the augmentation method increased the accuracy and 0 if it did not. Therefore, we model $T_i$ as a uniformly distributed binary random variable ($\mu_{T_i} = 0.5$, $\sigma_{T_i} = 0.5$) when assuming that the null hypothesis is true. We introduce the random variable $S$ as the sum of multiple $T_i$: $S = \sum_{i=1}^{N} T_i$. $S$ is distributed as a binomial distribution with $p = 0.5$, $n = N$. Finally, we let $s_e$ denote the empirically obtained sum of tests. The p-value, which is the probability of obtaining a result at least as extreme as the observed one under the null hypothesis, is given by

$$P(S \geq s_e | H_0) = \sum_{k=s_e}^{N} \binom{N}{k} 0.5^N.$$

Using the test accuracies from the experiments, we compute this probability, resulting in a p-value of 2%. This low p-value indicates that the observed improvement in test accuracy with Brownian motion augmentation is statistically significant, allowing us to reject the null hypothesis with considerable confidence.

### 1.3 Stretch-equivalent parameter estimation

This subsection details the calculation of the equivalent uniform stretch parameter of the Brownian motion data augmentation ($\sigma_e$). This augmentation method alters the event lengths, and our objective is to determine the relative standard deviation of the length variation. This parameter helps verify that uniformly stretching the data using $\sigma_e$ does not increase accuracy, but using the Brownian motion data augmentation does.

In the methods section, we introduced the trace array $X$. Here, we define the function $l(X) : \mathbb{R}^L \to \mathbb{N}$, which estimates the length of the event. This function employs a straightforward approach using two threshold levels ($h_s$ and $h_e$). The start of the event, $i_s$, is identified as the first instance when the trace falls below the start threshold $h_s$. The end of the event, $i_e$, is the last index where the trace value is below the end threshold $h_e$. Mathematically this can be expressed as

$$i_s(X) = min\ \{i : X[i] < h_s\},$$

$$i_e(X) = max\ \{i : X[i] < h_e\}.$$

And the event length is defined as

$$l(X) = i_e(X) - i_s(X).$$

While various length estimators could be proposed, our code demonstrates that this estimation is accurate for numerous samples. The thresholds used were $h_e = -0.25$ and $h_e = -0.5$

We denote the $i$th trace in our database as $X_i$ and its Brownian motion augmented version as $Y_i$, with $N_t$ being the total number of traces. To obtain the stretch equivalent parameter $\sigma e$, first we calculate the mean stretch factor $\mu_e$

$$\mu_e = \frac{1}{N_t} \sum_{i=1}^{N_t} \frac{l(X_i)}{l(Y_i)}.$$

Then, the variance $\sigma_e^2$ is

$$\sigma_e^2 = \frac{1}{N_t - 1} \sum_{i=1}^{N_t} \left( \frac{l(X_i)}{l(Y_i)} - \mu_e \right)^2 .$$

After using these formulas in our dataset augmented with Brownian motion of $\sigma = 0.9$, we obtained that the equivalent stretch was $\sigma_e \simeq 0.05$.

#### 1.4 YupanaNet tuning

As described in the methods section, the tuning of YupanaNet parameters was conducted without using any samples from the test dataset originally employed for QuipuNet. The original "train" dataset was split into training, validation, and test sets with 80%, 10% and 10% of the samples randomly selected, respectively. When discussing parameter tuning, the term "test set" refers to the samples from the original training dataset that were not used in the tuning trainings.

During each run for each neural network setting, the loss curves for the training and validation datasets were saved. Additionally, the accuracy results for the training, validation, and test datasets were recorded. In our analysis, we used the mean and standard deviation of all these results to compare different networks and select the optimal architecture for YupanaNet. The Adam optimizer [1] was utilized to fit the model.

Our initial model for tuning was QuipuNet with skip connections since QuipuNet was a state-of-the-art neural network, and extensive literature suggests that residual connections have potential accuracy improvements. We re-evaluated the parameters for augmentation steps, resulting in minor adjustments to the starting parameters for uniform stretching, noise addition, and magnitude multiplication.

Following the steps recommended by Yu and Zhu [2], we began by tuning the learning rate and subsequently selecting the batch size based on previous experiments. Next, we modified the architecture parameters. We first tuned the number of blocks and kernel sizes, testing several configurations with more blocks than QuipuNet. Then, we adjusted the number of units in the final dense layers. We also experimented with changing the activation functions from ReLUs to swish, but this did not result in any improvement. Following this, we compared different augmentation parameters, including the only parameter of the Brownian motion data augmentation, to fine-tune their performance. Finally, we introduced the attention layer to enhance the network further.

[2] Tong Yu and Hong Zhu. Hyper-parameter optimization: A review of algorithms and applications. *arXiv preprint arXiv:2003.05689*, 2020.
